# High-throughput drug screening for inhibition of influenza A virus infection based on human SIRT1 promoter and Genipin suppressing influenza A virus by activation of AMPK-SIRT1-PGC-1α signaling pathway

**DOI:** 10.1101/2024.07.10.602919

**Authors:** Jinghan Ye, Dekun Liu, Qianwen Wang, Jianping Dai

## Abstract

The energy metabolism crisis is considered an important risk factor for severe influenza A virus (IAV) infection. During virus replication, the host cell’s “metabolic reprogramming” is beneficial for increasing the energy demand of the virus. SIRT1 plays a major role in altering metabolic reprogramming, and upregulation of SIRT1 expression can defend against viral infection. This study established a high-throughput drug screening method for human SIRT1 promoter. Nine natural medicines were selected from 134 traditional Chinese medicines. Among them, the activity of *Gardenia jasminoides* Ellis was relatively high. Further research has found that the plant extract and its active compound Genipin and its derivatives can significantly inhibit IAV replication, improve the survival rate of infected mice, and inhibit pneumonia. In addition, Genipin significantly increased the levels of energy metabolism core regulatory factors SIRT1, PPAR γ, PGC-1 α, and p-AMPK, inhibited IAV induced activation of MAPKs and NF-κB, and alleviated inflammatory response. The pharmacological antagonists of SIRT1 and PGC-1 α, as well as siRNA, significantly counteracted the effects of Genipin on IAV replication and inflammation. In summary, we found that Genipin and its derivatives could significantly inhibit IAV replication and inflammation, possibly by activating the AMPK-SIRT1-PGC-1α signaling pathway and altering metabolic reprogramming.

## Introduction

In recent years, it has been noted that biochemical and energy metabolism play important roles in acute viral infections (1, 2). During the replication process, viruses must utilize a large number of macromolecules produced by host cells to synthesize their own proteins and nucleic acids. Especially for the budding of envelope viruses, it is necessary to consume a large amount of host fatty acids. In addition, the characteristic of severe acute influenza A virus (IAV) infection is a “cytokine storm”. Cytokine storm is also a high-energy intensive process (3). These processes require host cells to undergo metabolic reprogramming to support the virus’s higher demands for metabolic intermediates and energy. The host cell metabolism transitions to the Warburg effect (narrowly defined as aerobic glycolysis) (1, 2). At present, the “energy metabolism crisis” is considered an important risk factor for severe IAV infection (4), and a new and promising anti IAV strategy has been proposed to address “energy metabolism crisis” (5, 6).

Human sirtuins (SIRT1-7) are evolutionarily conserved NAD^+^-dependent deacetylases and ADP ribosyltransferases that regulate many basic cellular processes, including gene silencing, DNA repair, and metabolic regulation (7, 8). SIRT1 works in synergy with PPARs, AMPK, and PGC-1 α, participating in biochemical and energy metabolism in viral infections (9–11), inhibiting PKM2 which controls the Warburg effect (aerobic glycolysis) (12–14), and altering host cell “metabolic reprogramming” (15–21). There are reports that SIRT1 is a broad-spectrum antiviral host protein that serves as the core defense mechanism for hosts against DNA and RNA viral infections (22, 23). SIRT1 inhibitor EX-527 and siRNA SIRT1 can both increase the production of IAV offspring, while SIRT1 activators such as resveratrol, oligosaccharides, and CAY10602 can significantly promote the expression of SIRT1 *in vitro* and *in vivo*, inhibiting the replication of IAV (PR8) (23–25). In addition, SIRT1 activators can also inhibit the replication of many other viruses, such as hepatitis C virus (HCV), respiratory syncytial virus (RSV), vesicular stomatitis virus (VSV), human t-cell leukemia virus type 1 (HTLV-1), measles virus (MV), etc. (26–31). Pharmacological inhibition or silencing of SIRT1 by siRNA can promote the replication and spread of these viruses (26–31). In addition to energy metabolism, the SIRT1 signaling pathway also regulates the body’s oxidative levels and inflammatory responses through key transcription factors such as acetylated NF-κB and Nrf2 (32, 33). Therefore, SIRT1 is a good target for screening antiviral drugs, including IAV.

In China, many traditional Chinese medicine (TCM) have been used for thousands of years to treat IAV infections (34). In this study, we established a high-throughput drug screening method based on the human SIRT1 promoter and screened 134 TCM samples. We found that several TCMs could significantly stimulate the transcription of the SIRT1 promoter. Among them, *Gardenia jasminoides* Ellis had higher activity. In addition, we found that the crude extract and main active ingredients of this plant, Genipin and its derivatives (Geniposide, Genipin 1-β-D-gentiobioside and Shanziside), could significantly inhibit IAV infection. Finally, we focused on studying the anti-IAV mechanism of Genipin on SIRT1 related signaling pathways.

In addition, as a pulmonary fibrosis and depression inhibitor, Genipin has been applied for a patent in China, and several traditional Chinese patent medicines made from this compound have been marketed, but the anti-IAV activity of this compound has not yet been reported. Therefore, this study conducted a more in-depth study on the anti-influenza efficacy and antiviral mechanism of this compound, providing a useful basis for the promotion and application of these traditional Chinese patent medicines.

## Materials and methods

### Compounds and reagents

Medicinal plants were acquired from the Puning Pharmaceutical Market in Guangdong, China. The specimens are kept in our laboratory. Genipin and its derivatives (Geniposide, Genipin 1-β-D-gentiobioside and Shanziside) were purchased from MedChemexpress (MCE) Co., Ltd. (New Jersey, USA). TPCK trypsin (4370285), ribavirin (R9644), MTT, and DMSO were purchased from Sigma Aldrich, Inc. (St. Louis, MO, USA). Lipofectamine 2000 reagent, Pfu DNA polymerase, DNase, and TRIzzol reagent were purchased from Invitrogen Life Technologies. Inc. (Carlsbad, CA, USA). Resveratrol (510360) was acquired from Solarbio (Beijing, China). The luciferase detection kit was bought from Promega (Madison, WI, USA). Anti sirt1 (8469) antibody was purchased from Abcam (Cambridge, UK). ERK1/2 (8867), p-ERK1/2 (13148), p-JNK (3708), JNK (4671), p-p38 (4092), p38 (14451), NF - κ B p65, lamin B1, β - actin (12262) and secondary horseradish peroxidase coupled anti-rabbit and anti-mouse antibodies were bought from Cell Signaling Technology ® Inc. (Danvers, MA, USA). All other chemicals and solvents were commercially available and analytical grade.

### Construction of plasmids

To construct a SIRT1 promoter luciferase reporter plasmid, A549 cell genomic DNA was served as a template to amplify the human SIRTI (GeneBank: NC: 00010.11), we cloned the PCR product into the pGL3-basic vector, named pSIRT1-luc. We also cloned the human SIRTI gene (GeneBank: AF083106.2) into the pcDNA3.1 vector. Finally, we performed DNA sequencing validation on all plasmids. The primers were listed in **Table S1**.

### Cell, virus, and cytotoxicity tests

The cells employed in the experiment were A549 lung cancer cells and Madin Darby canine kidney (MDCK) cells. Both A549 and MDCK cells were cultured in Dulbecco modified Eagle medium (DMEM) containing 10% fetal bovine serum (Invitrogen, Carlsbad, CA, USA), and cultivated in a 5% CO_2_ humidified incubator. The virus stock was amplified using MDCK cells or 11 day chicken embryos, and the influenza virus strain used was A/Puerto Rico/8/34 (PR8, H1N1). The virus titer was determined through plaque formation assay (35). The cytotoxicity of all medicinal plant extracts, positive control drugs resveratrol, Genipin, and their derivatives was measured using MTT assay (36). Calculate the experimental drug concentration required to reduce cell viability by 50% (IC_50_). In our experiment, the maximum concentration without cytotoxicity was used as the drug treatment concentration for subsequent experiments. All IAV experiments were conducted in a Biosafety Level 2 (BSL-2) laboratory.

### High throughput drug initial screening test

The cell line used was A549 cells. A549 cells were inoculated into a 96 well microplate for 24 hours and allowed to grow to 80%. Then, the pSIRT1 luc and pRL-TK (internal control) plasmids were co-transfected using Lipofectamine 2000 reagent (Invitrogen). After 8 hours of cultivation, cells were infected with IAV PR8 virus strain (MOI=0.01) and treated with experimental drug dissolved in virus growth medium (VGM, containing MEM, 1ug/mL TPCK trypsin, and 0.125% (w/v) bovine serum albumin) for 24 hours. Changes in CPE were observed under an optical microscope at appropriate times. Measure luciferase activity using a dual luciferase assay kit (Promega, Madison, WI). And calculate the Z ’factor according to literature reports (37), a statistical parameter for quantifying the suitability of high-throughput drug screening models (37).

### In vitro antiviral test

Human lung cancer A549 cells were seeded on 6-well plates for 24 hours. Before the experiment, the virus was treated with continuously diluted test drug dissolved in VGM medium for 2 hours. Then wash A549 cells three times with PBS, add pre-treated virus solution (MOI=0.001), adsorb for 1 hour, and wash three times with PBS. Further incubate for 48 hours in the culture medium containing the test drug. After three cell freeze-thaw cycles, collect the supernatant, centrifuge, and determine the viral titer using the TCID_50_ method. Calculate 50% tissue culture infection dose unit (TCID_50_/mL) using Reed and Muench methods.

### Immunoblotting test

RIPA lysis buffer (Sigma) was used to extract total cell proteins, which contained phosphatase inhibitors and protease inhibitors. The cell samples were scraped with a cell scraper into a 1.5ml EP tube, boiled the sample at 100 ℃ for 10 minutes to denature. The protein concentration was determined using the BCA method (Thermo Scientific). Extract nuclear protein using EpiQuik nuclear protein extraction kit (Epigentek, Wuhan, China) to detect nuclear translocation of NF - κB p65. After adding 4 × SDS gel loading buffer to the sample, the micro sampler will load the sample with 25ul, and the sample will be subject to 10% SDS-PAGE electrophoresis. After transferring to the PDVF membrane, the membrane was blocked with 5% skim milk powder in TRIS buffered physiological saline. Then, incubated with first antibody (1:1000), shook gently on a shaking table overnight at 4 ℃. Subsequently, incubated with horseradish peroxidase coupled anti-rabbit or anti-mouse secondary antibodies. Finally, used the ECL detection kit (Thermo Fisher Scientific ™, Cleveland, OH, USA) to visualizes specific bands.

### Enzyme-linked immunosorbent assay

After 3 freeze-thaw cycles of cells, the samples of each experimental group were collected. After centrifuged 1500g for 10 minutes, the supernatants were collected and stored in a -80 ℃ ultra-low temperature freezer for analysis. Finally, use a commercialized ELISA detection kit to detect the levels of inflammatory cytokines (TNF-α, IL-6, IL-8, and IL-1 β), following the manufacturer’s instructions (Dakewe, Beijing, China).

### Real time quantitative PCR

Extract total RNA from cells and lung tissue using the Trizol RNA Purification Kit (Invitrogen). After diluting with TE solution appropriately, the absorption values at 260nm and 280nm were measured using a spectrophotometer to determine the concentration and purity of the RNA solution. Synthesize cDNA from 1 μg total RNA using an All-in-one cDNA Synthesis SuperMix Kit (biotool). The qRT-PCR reaction system (total volume 50 ul) included: 5 ul of cDNA, 0.5 ul of each upstream and downstream primers, 1 ul of Taq enzyme, 0.5 ul of dNTP, 10 ul of SYBR Green master mix, and 32.5 ul of ddH_2_O. The relative expression levels of each gene were expressed as 2^-ΔΔ^ ^Ct^. The primers used were listed in **Table S1**.

### SiRNA assay

Specific human SIRT1 siRNA, PGC-1 alpha siRNA, and control siRNA were purchased from Biomec Biotechnology Co., Ltd. (Nantong, Jiangsu, China). GoldenTran mRNA siRNA transfection reagents (Changchun Jinchuan Technology Co., Ltd.) were used to transfect SIRT1 siRNA, PGC-1 alpha siRNA, or control siRNA into A549 cells (1 × 10^6^). After 24 hours, the cells were infected (MOI=0.001) and treated with corresponding drugs at a temperature of 37 ℃ and a CO_2_ concentration of 5%. After 24 hours, TCID50 method was used to detect virus titer, and ELISA method was used to detect cytokine production (n=5).

### In vivo research

The experiment was conducted according to the ARRIVE guidelines. Animal experiments have obtained ethical approval from the Institutional Animal Care and Use Committee (IACUC) of Shantou University (Shantou, China) (No. SUMC 2022-033), March 26, 2022.

Half of male and female SPF Bal b/c mice aged 6-8 weeks were purchased from Beijing Charles River Laboratory Animal Technology Co., Ltd. (Beijing, China). The experimental animals were domesticated in an SPF level facility for 7 days, with a 12 h light and dark cycle, standard feed and water, and controlled environmental temperature and humidity.

The mice were randomly divided into 7 groups (n=16). Before infection, all mice were administered pentobarbital sodium 60mg/kg intraperitoneally for anesthesia. Blank control (BC) mice were intranasally instilled with 50 μ l of VGM culture medium and intraperitoneally injected with sterile physiological saline (0.5% DMSO) for 6 consecutive days (once a day). The mice in the negative control group (NC), positive control group (PC), and treatment group with Genipin and its derivatives were first infected with IAV 50 μl (PR8, 10 × MLD50) via nasal infection, and then injected intraperitoneally with sterile physiological saline (0.5% DMSO), ribavirin (50 mg/kg/d), Genipin and its derivatives (all 10 mg/kg/d) for 6 consecutive days.

Monitor and record the weight and survival rate of each group of mice daily for 14 consecutive days. On the 7th day, 6 mice were randomly selected from each group and euthanized under complete isoflurane anesthesia for cervical dislocation. The lung index is evaluated by measuring the percentage of lung wet weight (g) to body weight (g) (lung index=lung wet weight (g)/body weight (g) × 100%). The right lung of the mouse was fixed with 4% paraformaldehyde for pathological tissue sectioning. The left lung was frozen at -80 ℃ for the determination of various pathological and biochemical indicators. The expression of cytokines and viral load in lung tissue were detected using qRT PCR and TCID50 methods. Take the right lung and embed it in paraffin, stain it with hemoglobin and eosin (H&E), and observe the pathological changes at 200× and 400× magnification. Each slide was evaluated by two independent researchers using a blind method. The degree of pathological changes was evaluated using a semi-quantitative scoring system based on literature (38).

### Statistical methods

Statistical analysis was conducted using SPSS 19.0 software, using student tests or one-way ANOVA methods. All experimental data are expressed in mean ± SD. A *P*-value less than 0.05 is considered statistically significant.

## Results

### Establishment of drug screening methods and screening test results

Although many studies have shown that upregulation of SIRT1 can significantly inhibit IAV infection (23–25), in this study, we still first proved this conclusion. In **Fig. 1A**, transfection of SIRT1 overexpression plasmid did significantly inhibit IAV replication. Then, we constructed a human SIRT1 promoter luciferase reporter gene (pSIRT1 luc) with a length of 1844bp (**Fig. S1**). In addition, we used the SIRT1 agonist resveratrol (Res) to evaluate the feasibility of our drug screening method. The maximum non cytotoxic concentration of resveratrol we tested on A549 cells was 50 μ mol (**Fig. 1B** and **C**). Finally, we co-transfected pSIRT1 luc and pRL-TK plasmids into A549 cells to determine the effect of resveratrol on the transcriptional activity of the SIRT1 promoter. The results showed that the pSIRT1 luc reporter plasmid could transcribe normally without any stimulation (**Fig. 1D** and **E**), IAV infection could reduce the transcription of the SIRT1 promoter, but not significantly, while resveratrol could increase the transcription activity of the SIRT1 promoter (**Fig. 1D** and **E**). Finally, we calculated the Z ’factor for this study to be 0.58 (>0.5). According to the previously reported high-throughput drug screening criteria (37), our drug screening model was effective.

**Fig. 1.**
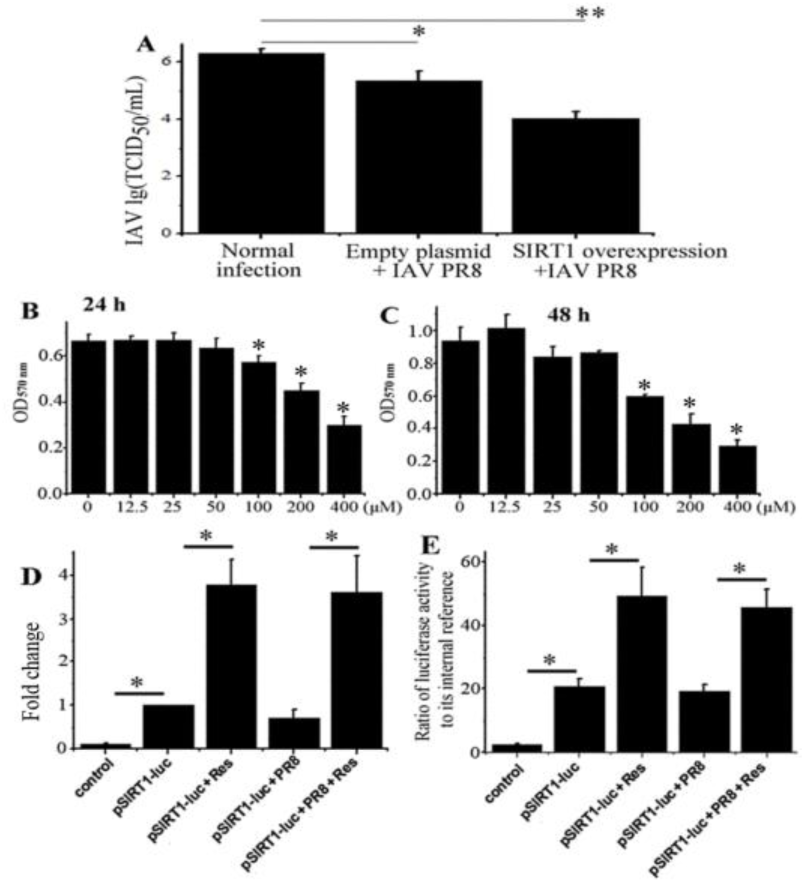
Feasibility evaluation of drug screening methods. (**A**) Overexpression of SIRT1 inhibited the replication of IAV. Firstly, A549 cells were transfected with pSIRT1 luc plasmid or control plasmid, respectively. After 8 hours, the cells were infected with IAV (PR8, Multiplicity of infection (MOI) = 0.01). Collect the supernatant 24 hours after infection (p.i.) and measure the viral titer using the TCID50 method. * *P* < 0.05, ** *P* < 0.01 vs the normal infection group. (**B** and **C**) Measure the cytotoxicity of resveratrol on A549 cells using MTT assay at 24 or 48 hours, respectively. * Compared with 0 μ mol, *P*<0.05. (**D** and **E**) The effect of resveratrol on the transcriptional activity of the SIRT1 promoter. The control group was co-transfected with empty vector (pGL3 basic) and pRL-TK plasmid (internal reference) into A549 cells. In the pSIRT1 luc and pSIRT1 luc + Res groups, A549 cells were first co-transfected with pSIRT1 luc and pRL-TK plasmids, and treated with 0.5% DMSO and 50 μmol resveratrol (Res), respectively. In the pSIRT1-luc + PR8 and pSIRT1-luc + PR8 + Res groups, after co-transfected for 8 h, PBS washing and IAV infection (MOI=0.01) were performed, and then treated with 0.5% DMSO and 50 μmol Res, respectively. After 24 hours, qPCR was used to detect the transcription level of firefly luciferase gene, and the result was expressed as 2^-ΔΔCt^. The luciferase activity was measured using the luciferase reporter gene detection kit (Promega), and the ratio of luciferase activity to its internal control in each treatment was calculated. The data were represented by the mean ± standard deviation of three independent experiments, * *P* < 0.05。

Based on this drug screening method, 134 traditional antiviral Chinese medicine samples were screened (**Table S2**), and it was found that the transcription activity of the SIRT1 promoter in 9 Chinese medicine samples increased by more than three times. These traditional Chinese medicines include *Citrus aurantium* L., *Gentiana macrophylla* Pall, *Angelica sinensis* (Oliv) Diels, *Gardenia jasminoides* Ellis, *Aster tataricus* L. f., *Centella asiatica* (L.) Urb., *Imperata cylindrica* Beauv. var. major (Nees) C. E. Hubb., *Vitex trifolia* L., and *Spirodela polyrrhiza* (L.) Schleid. Among them, the anti-IAV effects of *Citrus aurantium* L., *Gentiana macrophylla* Pall. and *Angelica sinensis* (Oliv.) Diels had been reported (39–41). After excluding traditional Chinese medicine with existing reports or unclear active ingredients, we chose *Gardenia jasminoides* Ellis as the drug of interest for further research.

### Genipin and its derivatives enhanced the activity of pSIRT1 luc promoter and inhibited IAV infection *in vitro*

On the basis of drug screening, the anti IAV activity of *Gardenia jasminoides* Ellis extract was tested. From **Fig. 2 A** and **B**, it could be seen that the extract of *Gardenia jasminoides* Ellis had no significant cytotoxicity within the concentration range of 12.5 μ g/mL to 100 μ g/mL, and could significantly inhibit IAV infection within the concentration range of 6.25 μ g/mL to 50 μ g/mL. Then, we investigated the effects of the main active components of this plant, Genipin and its derivatives (Genipin 1-β - d - geniobioside and Shanziside), on the transcriptional activity of the pSIRT1 luc promoter and IAV infection *in vitro* (**Fig. 2C**).

**Fig. 2.**
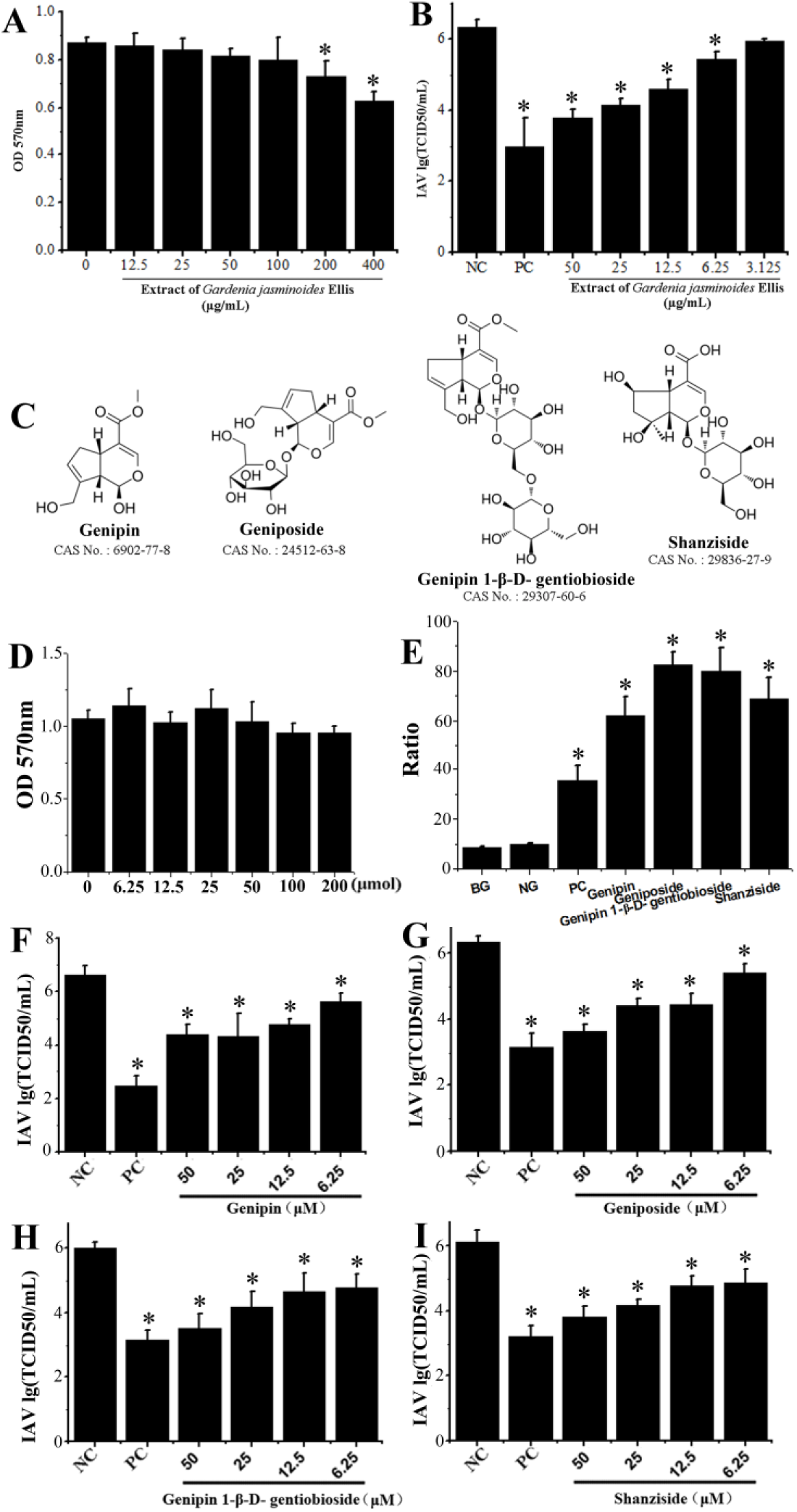
The effects of Genipin and its derivatives on the pSIRT1 luc reporter and IAV infection *in vitro*. (**A**) MTT assay was used to determine the cytotoxicity of *Gardenia jasminoides* Ellis extract on A549 cells. (**B**) The inhibitory effect of *Gardenia jasminoides* Ellis extract on IAV (PR8) infection *in vitro*. (**C**) The structure of Genipin and its derivatives. (**D**) MTT assay was used to determine the cytotoxicity of Genipin on A549 cells. (**E**) The stimulating effect of Genipin and its derivatives on the pSIRT1 luc reporter. All groups were co-transfected with pSIRT1 luc and pRL-TK plasmids. Except for the control group (BG), all other groups were infected with IAV (MOI=0.01). The negative group (NG) and positive control group (PC) were treated with 0.5% DMSO and 50 μmol resveratrol (Res), respectively. All other groups were given Genipin and its derivatives at a dose of 50 μ mol. The data is expressed as the ratio of luciferase activity to its internal control. (**F, G, H** and **I**) The inhibitory effect of Genipin and its derivatives on IAV (PR8) infection. The negative control (NC) A549 cells were only infected with IAV (MOI=0.01) and treated with DMSO (0.5%, v/v). In the PC, Genipin, and their derivatives groups, A549 cells infected with IAV were treated with ribavirin (20 μg/mL), Genipin, and Genipin derivatives (6.25, 12.5, 25, 50 μmol), respectively. After 48 hours, collect the supernatant and measure the titer using the TCID50 method. The data were the mean ± standard deviation of three independent experiments. * Compared with the NC group, P<0.05.

Initially, we also determined the cytotoxicity of these compounds on A549 cells. As shown in **Fig. 2D**, within the concentration range of 6.25 μ mol∼200 μ mol, Genipin showed no significant cytotoxicity on A549 cells. In addition, the derivatives of Genipin also showed no significant cytotoxicity on A549 cells within the concentration range of 6.25 μ mol to 200 μ mol (data not shown). As shown in **Fig. 2E**, Genipin and its derivatives could significantly stimulate the transcriptional activity of the SIRT1 promoter, with a ratio of 6.4573 ± 1.2288 in the Genipin group compared to the negative group. Then, through the TCID_50_ experiment, we found that Genipin and its derivatives could significantly inhibit IAV (PR8) infection in the concentration range of 6.25 μ mol to 50 μ mol (**Fig. 2 F, G, H**, and **I**).

### Genipin could inhibit IAV replication, suppress lung inflammation, and improve lung tissue pathological changes in the body

Subsequently, we evaluated the potential activity of Genipin against IAV infection *in vivo*. As shown in **Fig. 3**, Genipin could significantly improve the survival rate and average survival time of infected mice (**Fig. 3A**). On the 7th day, Genipin significantly reduced pulmonary virus replication, lung index (pulmonary edema), and production of lung cytokines IL-1 β, IL-6, and TNF - α (**Fig. 3 B, C**, and **D**). In addition, Genipin could improve the pathological changes of lung tissue induced by IAV, reduce alveolar exudation, alveolar wall destruction, and alveolar bleeding (**Fig. 4**).

**Fig. 3.**
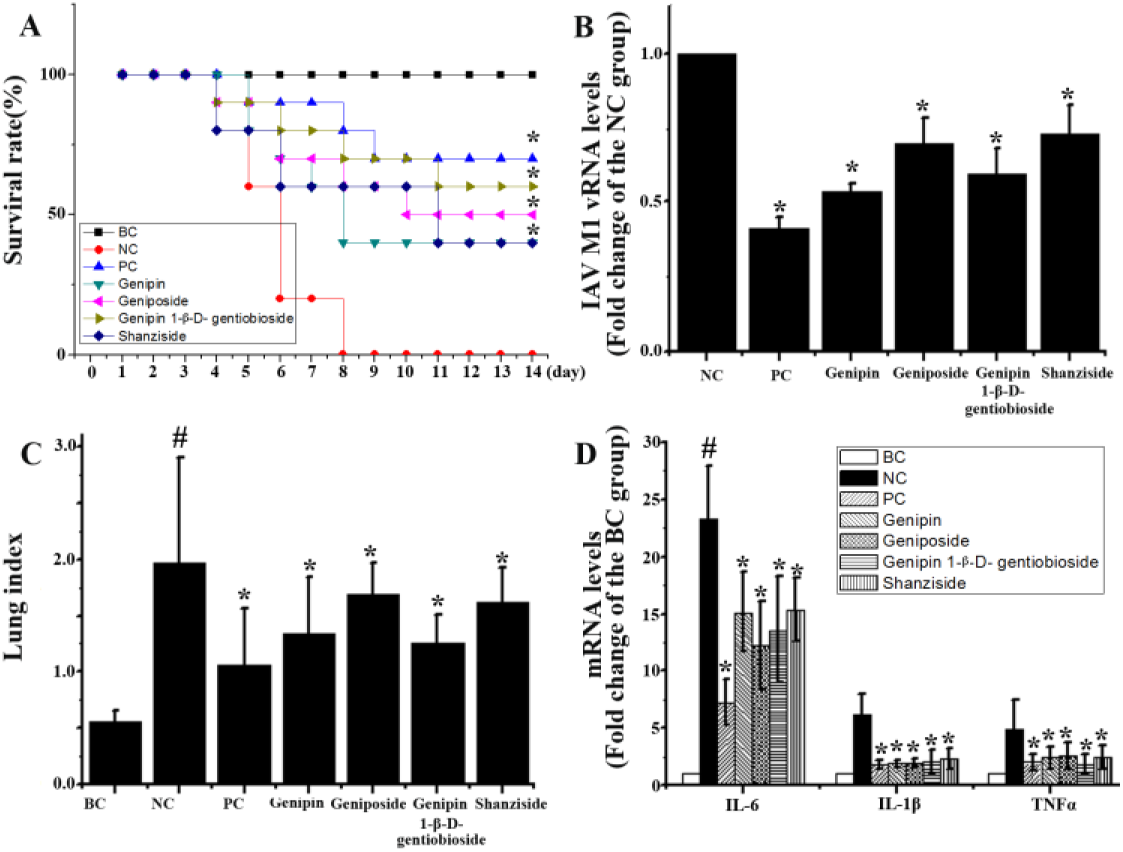
*In vivo* anti IAV activity of Genipin and its derivatives. In the blank control (BC), mice were intraperitoneally treated with VGM and gavage with sterile physiological saline. In the negative control group (NC), positive control group (PC), and the treatment groups with Genipin and its derivatives, the mice were infected with 10 × MLD50 IAV (PR8), were treated with sterile physiological saline (NC group), ribavirin (50 mg/kg/d, PC group), Genipin (10mg/kg/d) and its derivatives (10mg/kg/d), respectively. (A) Monitor the survival rate for 14 days. Kaplan-Meier analysis with Log-rank and Breslow test were used to analyze the significant differences in average survival time. (B) On the 7th day, the lung virus load was detected by qPCR. (C) Evaluate lung index by measuring the percentage of lung wet weight (g) to body weight (g) on the 7th day (lung index=lung wet weight (g)/body weight (g) × 100%). (D) On the 7th day, lung cytokine levels were detected by qPCR. The data shown are the mean ± SD. Survival rate was measured in 10 mice (n=10), while lung index, lung virus load, and lung cytokine levels were measured in 6 mice (n=6). Compared with the BC group, ^#^ P<0.05, and compared with the NC group, * P<0.05.

**Fig. 4.**
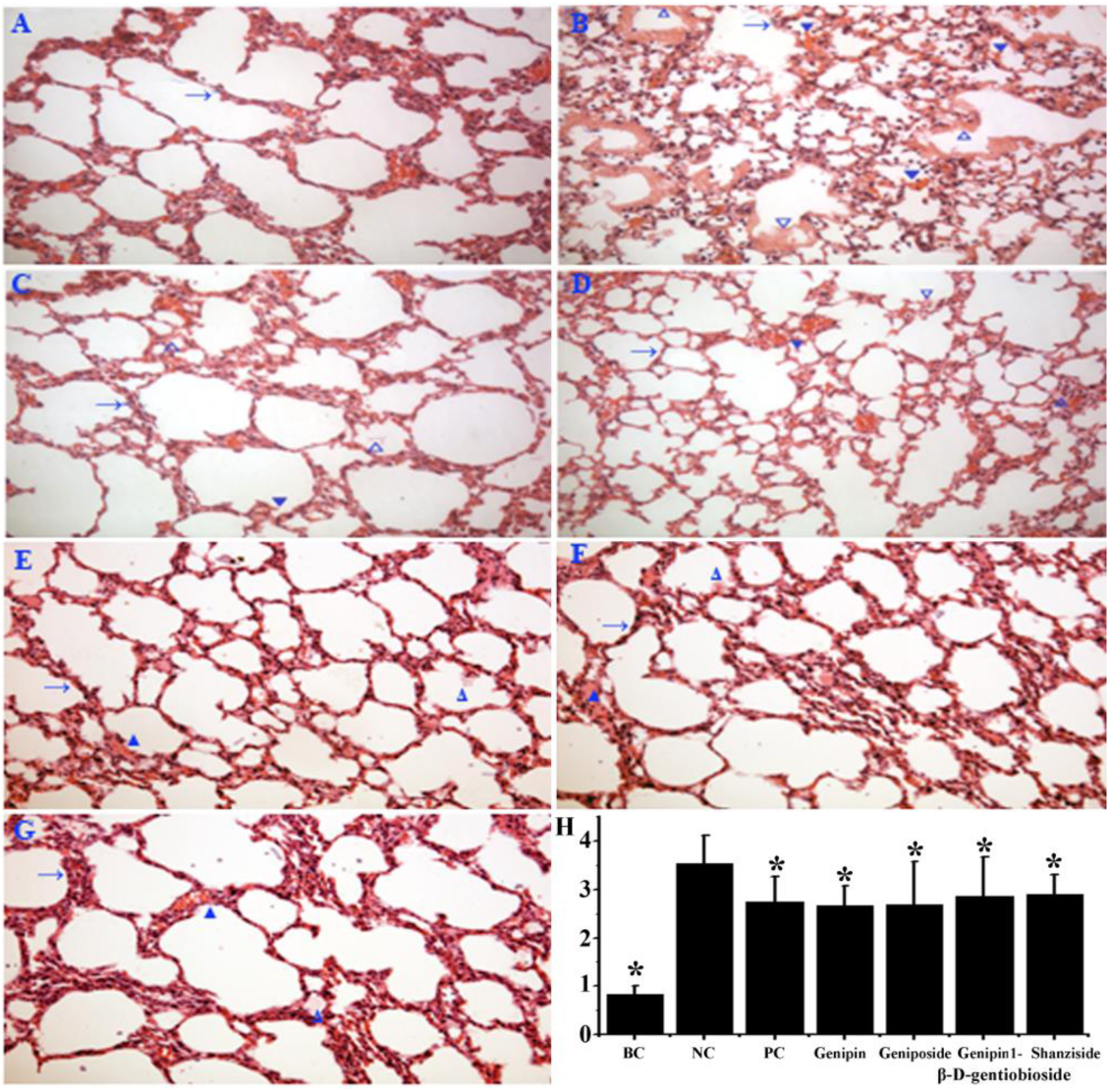
The effect of Genipin and its derivatives on pathological changes in mouse lung tissue. The treatment method for mice was shown in Figure 3. On the 7th day, 6 mice in each group were euthanized. Then perform H&E staining on the right lung. (A) the blank control group (BC), (B) the negative control group (NC), (C) the positive drug control group (PC), (D) Genipin treatment group, (E) Geniposide treatment group, (F) Genipin 1-β -D-gentiobioside treatment group, (G) Shanzisaide treatment group, (H) histopathological evaluation. (→) alveolar wall, (△) inflammatory exudation, (▼) bleeding. The magnification was 200 times.

### Genipin upregulates the expression of SIRT1, PPAR γ, and PGC-1 α *in vitro*, inhibiting the activation of IAV induced MAPKs and NF - κ B pathways

As shown in **Fig. 5**, IAV infection significantly downregulated the expression of SIRT1 and PGC-1 α (**Fig. 5A**), downregulated the phosphorylation of AMPK detected at 48 hours p.i (**Fig. 5D**), while Genipin significantly increased the expression of SIRT1, PPAR γ, and PGC-1 α (**Fig. 5A**), and upregulated the phosphorylation of AMPK (p-AMPK/AMPK) (**Fig. 5D**). IAV infection and Genipin treatment had almost no effect on the expression of AMPK (AMPK/β - actin) (**Fig. 5D**). Meanwhile, IAV infection significantly increased the phosphorylation of MAPKs (ERK, p38, and JNK), as well as the nuclear translocation of NF - κB p65, while Genipin significantly inhibited the phosphorylation of MAPKs (ERK, p38, and JNK) caused by IAV infection, as well as the overall expression and nuclear translocation of NF - κB p65 (**Fig. 5 B** and **C**). Moreover, the level of IAV viral NP protein was inhibited by drugs.

**Fig. 5.**
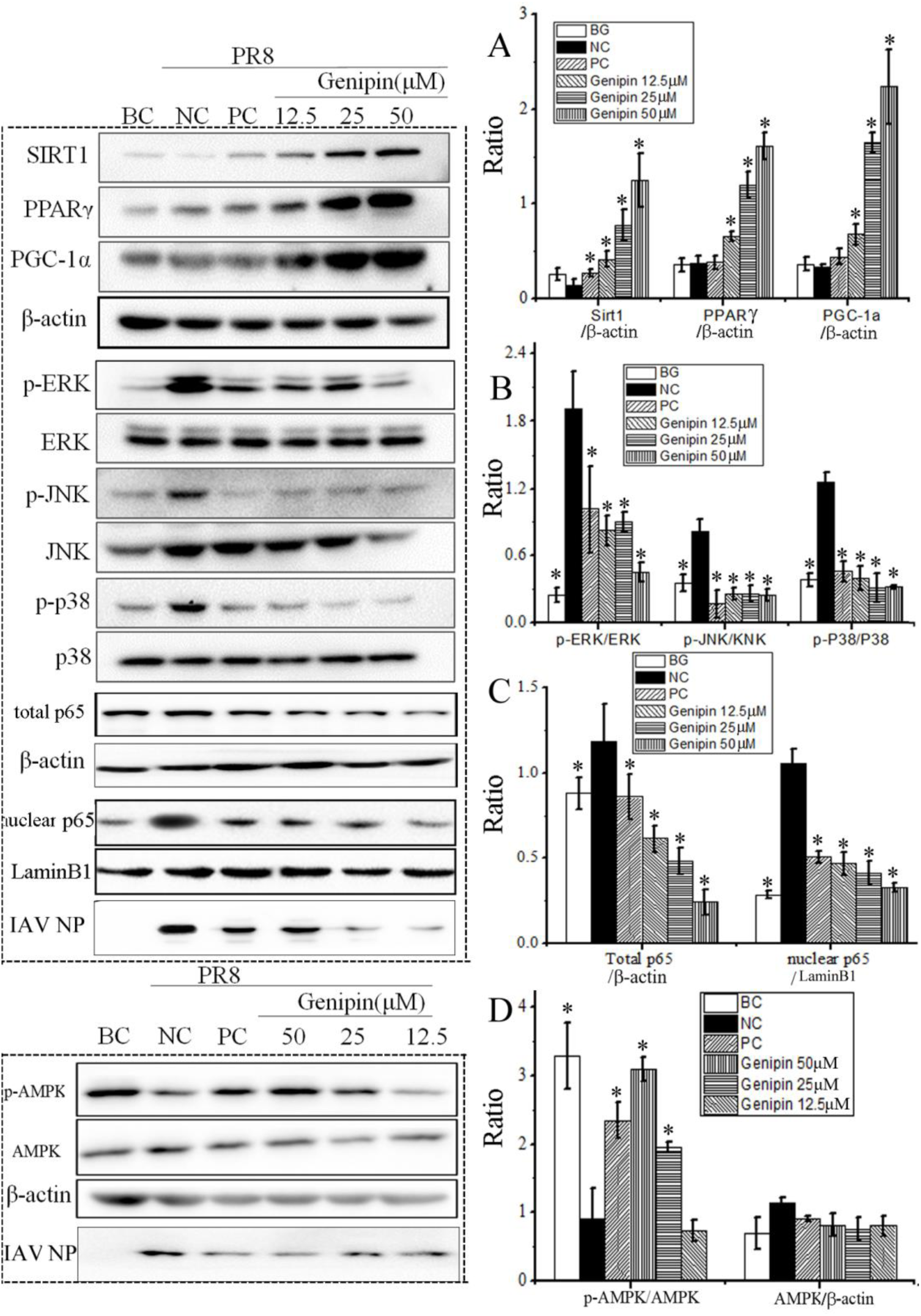
The effects of Genipin on PPAR γ, SIRT1, AMPK, PGC-1 α, MAPK (ERK, p38, and JNK), and NF - κ B signaling pathways after IAV infection *in vitro*. In the blank control (BC), A549 cells were treated with only 0.5% (v/v) DMSO. A549 cells in the negative control group (NC), positive drug control group (PC), and Genipin treated group were first infected with IAV (MOI=0.01), and then treated with 0.5% DMSO, ribavirin (20 μ g/mL), and Genipin (12.5 μ M, 25 μ M, 50 μ M), respectively. After 48 hours, the cell lysate was collected. Western blotting was used to detect the expression of SIRT1, PPAR γ, AMPK and PGC-1 α, the phosphorylation levels of MAPK (ERK, p38, JNK), AMPK, and total expression of NF -κB p65 and its nuclear translocation. The experiment was repeated 5 times (n=5), and compared with the NC group, * P<0.05.

### Genipin inhibits the expression of pro-inflammatory cytokines induced by IAV

As shown in **Fig. 6**, under the stimulation of IAV infection, the expression of pro-inflammatory cytokines (IL-1 β, IL-6, and TNF-α) significantly increased. But compared with NC control, Genipin significantly inhibited the expression of pro-inflammatory cytokines.

**Fig. 6.**
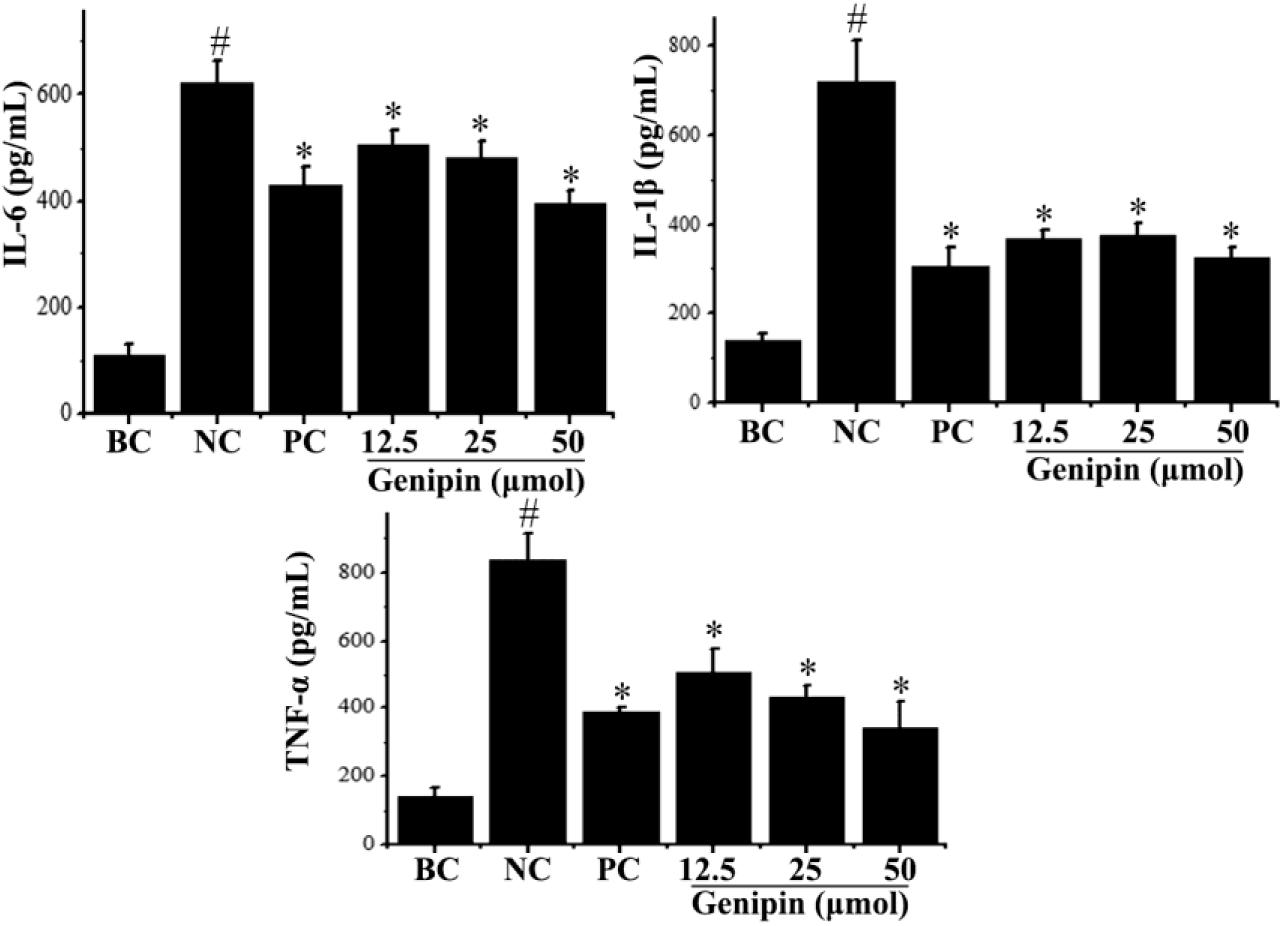
The effect of Genipin on cytokine production after IAV infection. In the blank control (BC), A549 cells were treated with only 0.5% (v/v) DMSO. A549 cells in the negative control group (NC), positive drug control group (PC), and Genipin treated group were first infected with IAV (MOI=0.01), and then treated with 0.5% DMSO, ribavirin (20 μ g/mL), and Genipin (12.5 μ M, 25 μ M, 50 μ M), respectively. After 48 hours, collect the cell supernatant for ELISA detection. The data were shown as the mean ± standard deviation of three independent experiments. ^#^ Compared with the BC group, *P*<0.05; * Compared with the NC group, *P*<0.05.

### Drug antagonists and siRNA of SIRT1 and PGC-1 α counteracted the effects of Genipin on IAV infection and pro-inflammatory cytokine expression

We further conducted *in vitro* antagonistic experiments on Genipin. As shown in **Fig. 7**, the SIRT1 antagonist nicotinamide (NAM) and PGC-1 α antagonist SR-18292 could significantly counteract the inhibitory effect of Genipin on IAV infection (**Fig. 7A**) and the inhibitory effect on IL-1 β and TNF - α production (**Fig. 7B**). SIRT1 siRNA and PGC-1 α siRNA also significantly counteracted the inhibitory effect of Genipin on IAV infection (**Fig. 7E**) and on IL-1 β and TNF - α production (**Fig. 7F**).

**Fig. 7.**
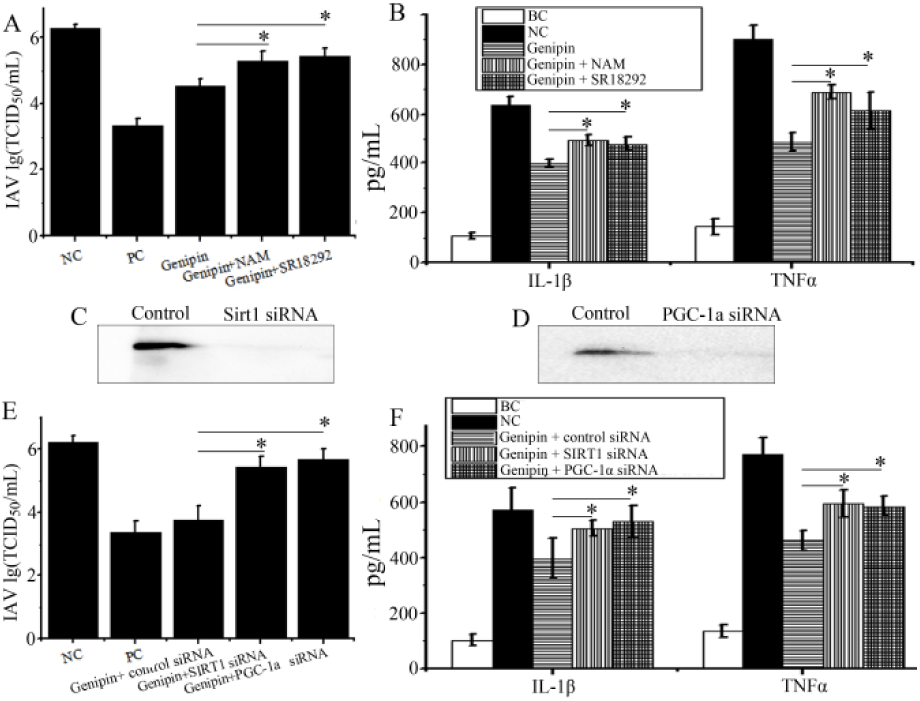
An antagonistic experiment on the effect of Genipin. The blank control group (BC), negative control group (NC), positive drug control group (PC), and genipin (12.5 μ mol) treatment group were the same as Fig. 5. In pharmacological antagonistic experiments, SIRT1 antagonist nicotinamide (NAM, 10mM) or PGC-1 alpha antagonist SR-18292 (15 μ M) were added together with Genipin. 48 hours later, TCID50 method was used to detect virus titer (**A**), and ELISA method was used to detect cytokine levels (**B**). In the siRNA antagonist assay, SIRT1 siRNA, PGC-1 α siRNA, or control siRNA were transfected with GoldenTran mRNA siRNA transfection reagents (Changchun Jinchuan Technology Co., Ltd.). After 24 hours, cells were infected with IAV (MOI=0.001) and treated with corresponding drugs. After 24 hours, Western blotting was used to determine the knockout effect of siRNA (**C** and **D**), TCID50 method was used to measure virus titer (**E**), and ELISA method was used to measure cytokine production (**F**). The data shown were the mean ± standard deviation of three independent experiments. Compared with the Genipin treatment group, \**P*<0.05.

## Discussion

SIRT1 is a broad-spectrum, evolutionarily conserved antiviral protein (26–31) that synergizes with PPARs, AMPK, and PGC-1 α, playing important roles in biochemical and energy metabolism (11, 42). PPARs can stimulate the expression of SIRT1 (43). AMPK can promote the synthesis of NAD^+^ required for intracellular SIRT1 activity (44). SIRT1 can serve as a co-activator of PPAR alpha complexes and activate PGC-1 alpha (45, 46), playing an important role in mitochondrial biogenesis and energy metabolism (47). In addition, there are reports that upregulation of SIRT1 expression can inhibit the replication of IAV, while siRNA mediated knockdown of SIRT1 can increase the production of IAV offspring (23–25). In addition to SIRT1, PPAR - α/γ and AMPK agonists can also provide protection against highly pathogenic influenza viruses in mice and influenza pneumonia patients (48, 49). Therefore, in this study, we established an antiviral drug screening model based on the human SIRT1 promoter luciferase reporter.

In this study, we screened 134 traditional Chinese medicines using pSIRT1 luc reporter and obtained 9 medicinal plants that significantly increased SIRTI expression. Among them, *Gardenia jasminoides* Ellis had high activity. In addition, we also explored the anti IAV effects of the active compounds (Genipin and its derivatives) of this plant. In *in vitro* studies, we have demonstrated that Genipin and its derivatives (Genipin-1-β - d-gentiobioside and Shanziside) can significantly inhibit the replication of IAV (PR8). In *in vivo* studies, we have demonstrated that these compounds can significantly improve the survival rate of infected mice, inhibit pulmonary virus titers, inflammatory factors, lung index, and pathological changes in lung tissue.

We found that the mechanism of action of Genipin against IAV might be to increase the expression of PPAR γ and PGC-1 α, and the phosphorylation of AMPK *in vitro*. In addition, it has been demonstrated that the MAPK (ERK, P38 and JNK ) and NF - κ B signaling pathways play important roles in influenza virus replication, and activating these pathways is beneficial for IAV infection(50–52). Inhibiting p38 MAPK activation can inhibit IAV replication, vRNP output, and cell apoptosis(50). Inhibiting ERK activation can block the nuclear entry and nuclear output of IAV vRNP (51). Inhibiting JNK and NF - κB activation can impair the synthesis of IAV vRNA and reduce the expression level of pro-inflammatory factors (52). The excessive activation of these signaling pathways can also induce cytokine storms, which are important pathological factors in IAV related ALI and ARDS (53). In addition, activation of SIRT1, PPAR γ, or PGC-1 α can inhibit the activation of the p38/JNK/ERK MAPKs pathway and NF - κ B pathway, suppressing inflammation (54–57). SIRT1 directly inhibits NF - κB signal transduction by deacetylating the NF - κ B p65 subunit. Overexpression of SIRT1 enhances the binding of PPAR α to RelA/p65 and induces its deacetylation (58). Conversely, the promoter of the SIRT1 gene contains many putative binding sites for NF - κB transcription factors, and activation of NF - κ B can downregulate the activity of SIRT1 (57).

In this study, we found that Genipin could significantly inhibit the activation of IAV induced MAPK (ERK, JNK, p38) and NF - κB signaling pathways, and further significantly reduce the expression of pro-inflammatory factors (IL-1 β, IL-6, and TNF – α) after IAV infection. In fact, many other studies have also shown that Genipin has anti-inflammatory, anti-infective, and antioxidant properties (59–61).

Moreover, pharmacological antagonists and siRNAs of SIRT1 and PGC-1 α could significantly counteract the effects of genipin on IAV replication and the production of IL-1 β and TNF - α. The entire experimental design and drug action mechanism are shown in **Fig. 8**.

**Fig. 8.**
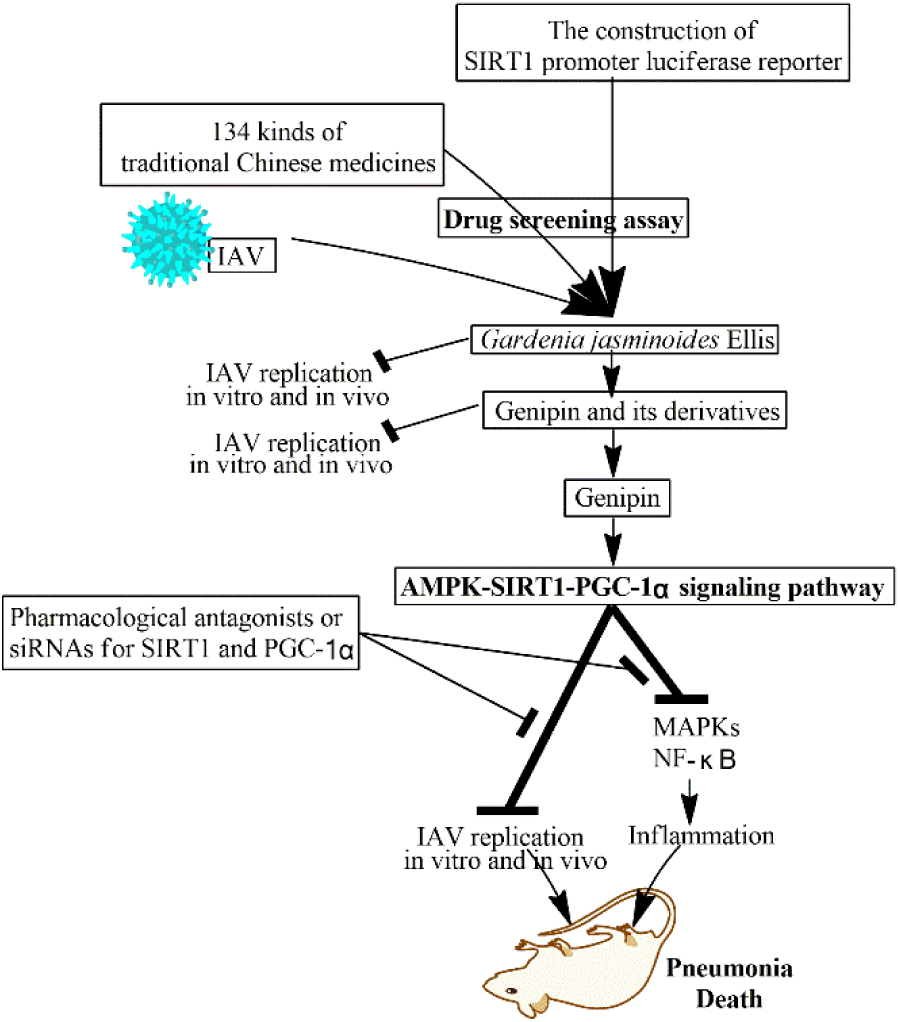
The whole experimental design and drug action mechanism

## Conclusion

Through an antiviral drug screening model based on the human SIRT1 promoter luciferase reporter, we obtained 9 traditional Chinese medicine samples with high transcriptional activity of the SIRT1 gene. *Gardenia jasminoides* Ellis and its main active components, Genipin and Genipin derivatives could significantly inhibit the replication of IAV (PR8) *in vitro* and *in vivo*. The anti IAV mechanism of Genipin might be to activate the AMPK-SIRT1-PGC-1 alpha signaling pathway, thereby inhibiting the activation of MAPK and NF - κ B pathways.

## SUPPLEMENTARY INFORMATION

The online version contains supplementary material available at

## AUTHOR CONTRIBUTIONS

Jinghan Ye, Qianwen Wang and Dekun Liu contributed to experimental data collection, analysis, and investigation. Qianwen Wang contributed to experimental data collection, Jianping Dai conceived and designed the study, performed the literature review and drafted the manuscript. All authors approved the final manuscript and agreed to take responsibility for all aspects of the work.

## Data availability

All the data are contained within the manuscript and supplementary Materials.

## DECLARATIONS

### Ethics Approval

This study was conducted in accordance with the Declaration of Helsinki. The animal experiment was conducted in accordance with China’s animal welfare legislation, and an ethical approval was obtained from the Institutional Animal Care and Use Committee (IACUC) of Shantou University (Shantou, China) (No. SUMC 2022-033) on 26 March 2022

### Consent for Publication

Not applicable.

